# The Neuroactive Potential of the Elderly Human Gut Microbiome is Associated with Mental Health Status

**DOI:** 10.1101/2024.08.08.607034

**Authors:** Paulina Calderón-Romero, Benjamin Valderrama, Thomaz Bastiaanssen, Patricia Lillo, Daniela Thumala, Gerard Clarke, John F Cryan, Andrea Slachevsky, Christian González-Billault, Felipe A. Court

## Abstract

Ageing is usually associated with physiological decline, increased mental health issues, and cognitive deterioration, alongside specific changes in the gut microbiome. However, the relationship between the neuroactive potential of the gut microbiome and mental health and cognition among the elderly remains less explored. This study examines a cohort of 153 older Chilean adults with cognitive complaints, assessing anthropometric data, mental health via five distinct tests, and gut microbiome composition through 16SV4 sequencing. Our findings reveal associations between anthropometric factors and depression scores in mental tests of participants with their gut microbiome composition. Notably, depression was associated with changes in the abundance of *Lachnospiraceae Eubacterium xylanophilum group* and *Fusobacteriaceae Fusobacterium*. Additionally, bacterial pathways involved in metabolising neuroactive compounds such as tryptophan, short-chain fatty acids, p-cresol, glutamate, and nitric oxide were associated with participant age, sex, and cognitive performance. Moreover, participants’ sex was associated with the neuroactive potential of specific bacteria, suggesting a role of the gut microbiome in sex-related mental health differences in the elderly. Together, to the best of our knowledge, this study demonstrates for the first time the association between the neuroactive potential of the human gut microbiome and mental health status in older individuals with cognitive complaints.

## 1. Introduction

According to global projections, the percentage of people over 60 years old will nearly double by 2050 [1]. As the global population ages, health concerns related to ageing are becoming increasingly important. Among the conditions affecting older adults, physical diseases, as well as cognitive disorders such as dementia and depression are the most common [2–4]. Ageing is a natural and complex process characterised by a steady decline in mental sharpness and independence in daily activities, the extent of which can be determinant for ageing well and healthy [5,6]. However, the mechanisms underlying the decline and the factors driving its individual-specific differences in the rate of decline remain to be fully elucidated.

Within the spectrum spanning from healthy ageing to dementia, cognitive complaints defined as self- or hetero-reported experiences of memory loss or cognitive decline—have emerged as a significant risk factor for future cognitive impairment, functional decline, and progression to dementia [7]. These complaints have been linked to varying degrees of cognitive and functional impairment, as well as the presence of neuropsychiatric symptoms [8]. Identifying individuals within the community who report cognitive disturbances and are at higher risk for functional decline and dementia conversion could pave the way for more personalised prevention strategies tailored to this high-risk group [9]. In this context, specific mental tests can be utilised to assess changes in brain function during ageing and to stratify risk groups. Recent improvements in test sensitivity and specificity have increased their accuracy in assessing cognitive issues in older individuals. Several tests have been developed and validated to cover different aspects of mental health in older people, including tests such as Activities of Daily Living Questionnaire (ADLQ) [10], Everyday Cognition (ECog) [11], Alzheimer Disease 8 (AD8) [12,13], Cognitive Reserve Scale (CRS) [14] and the Geriatric Depression Scale (GDS) test [15,16]. These provide physicians with valuable information for classifying cognitive level, memory loss, and other cognitive complaints that could affect daily life.

Along with well-established potential decline in brain functions and mental health, new studies show evidence supporting alterations in the gut microbiota of people transitioning from adulthood to elderly. These changes are characterised by a loss of overall microbial diversity and a greater inter-individual variation, also referred to as a higher degree of gut microbiome uniqueness [17,18]. Additionally, evidence suggests ageing is characterised by a decrease in Firmicutes and an increase in Bacteroidetes phyla [19], and that longevity has been associated with an increased abundance of *Clostridium cluster XIV*, Ruminococcaceae, *Akkermansia* and Christensenellaceae, known to produce molecules that may modulate human inflammation [20–24]. Moreover, it’s been suggested that age-related changes in the gut microbiome can accelerate the progression of ageing and inflammation, impacting overall well-being [25]. Therefore, changes in the gut microbiome have recently gained interest as factors that could distinguish between healthy and unhealthy ageing in the human population. Indeed, research has suggested a role for the gut microbiome as a potential mediator of overall health in older populations [17,26–28].

Interestingly, the gut microbiome has also been described as a modulator of the bidirectional communication between the brain and the gut, leading to the coining of the term microbiome-gut-brain axis [29]. Evidence suggests that gut bacteria can produce neuroactive metabolites [30,31], such as short chain fatty acids (SCFAs; acetate, propionate, butyrate) [32], which may modulate neuronal inflammation and has been linked to depression status [33]. Therefore, targeting the gut microbiome could lead to an improvement in mental health of the elderly, as depression is a common feature. Indeed, patients with major depressive disorder experience a reduction in SCFAs, while supplementation with butyrate has shown antidepressant effects in animal studies, improving intestinal permeability [34,35]. Moreover, depression is often associated with a loss of microbial stability in patients, which has been hypothesised to exacerbate depressive symptoms[36]. On the other hand, the production of bile acids and neurotransmitters may lessen the severity of depression [37]. Bacterial genera, such as *Enterobacter* and *Burkholderia* have shown a positive correlation with depressive symptoms and negative correlations with brain structures responsible for memory and emotional regulation [38].

In addition to SCFAs, other gut bacteria-derived metabolites may interact with the vagus nerve, the main direct route of communication between the central and peripheral nervous systems [39]. Therefore, the microbiota-gut-brain axis plays crucial roles in various processes impacting human mental health, including neurodevelopment, psychological and psychiatric behaviours, age-associated disorders, and neurodegenerative diseases [40,41]. However, whether the gut microbiome plays a role in modulating the mental health of the elderly, or its involvement in the decline in mental capabilities is not fully understood.

In this context, our study provides insights on how the composition and neuroactive potential of the gut microbiome are linked to cognitive performance, depressive symptoms, and other physiological factors. To the best of our knowledge, this is the first-time that changes in the neuroactive potential of the human gut microbiome -determined based on its inferred genetic content-have been linked to performance changes in cognitive assessments in a cohort of older individuals with cognitive dysfunction. Therefore, our findings offer new insights into the role of the gut microbiota as a modulator of mental health of this population and represents an initial exploration on the potential of the gut microbiome as a target for therapeutic interventions in the context of mental health in the elderly.

## 2. Results

### 2.1 The gut microbiome composition of the GERO cohort is primarily influenced by anthropometric variables and depressive symptoms scores

16S amplicon sequencing was used to characterise the taxonomic composition of the gut bacterial microbiome in the cohort. Results indicate that the microbiome of the GERO cohort is predominantly composed of the phyla Firmicutes (Mean = 53.7%, SD = 13.9%) and Bacteroidota (Mean = 34.1%, SD = 14.8%) **(Figure 1A)**. Nine additional human-commensal phyla were detected in much lower abundance, including Proteobacteria, Verrucomicrobiota, Actinobacteriota, Synergistota, Desulfobacterota, Campylobacterota, Elusimicrobiota, Deferribacterota, and Fusobacteriota.

**Figure 1.**
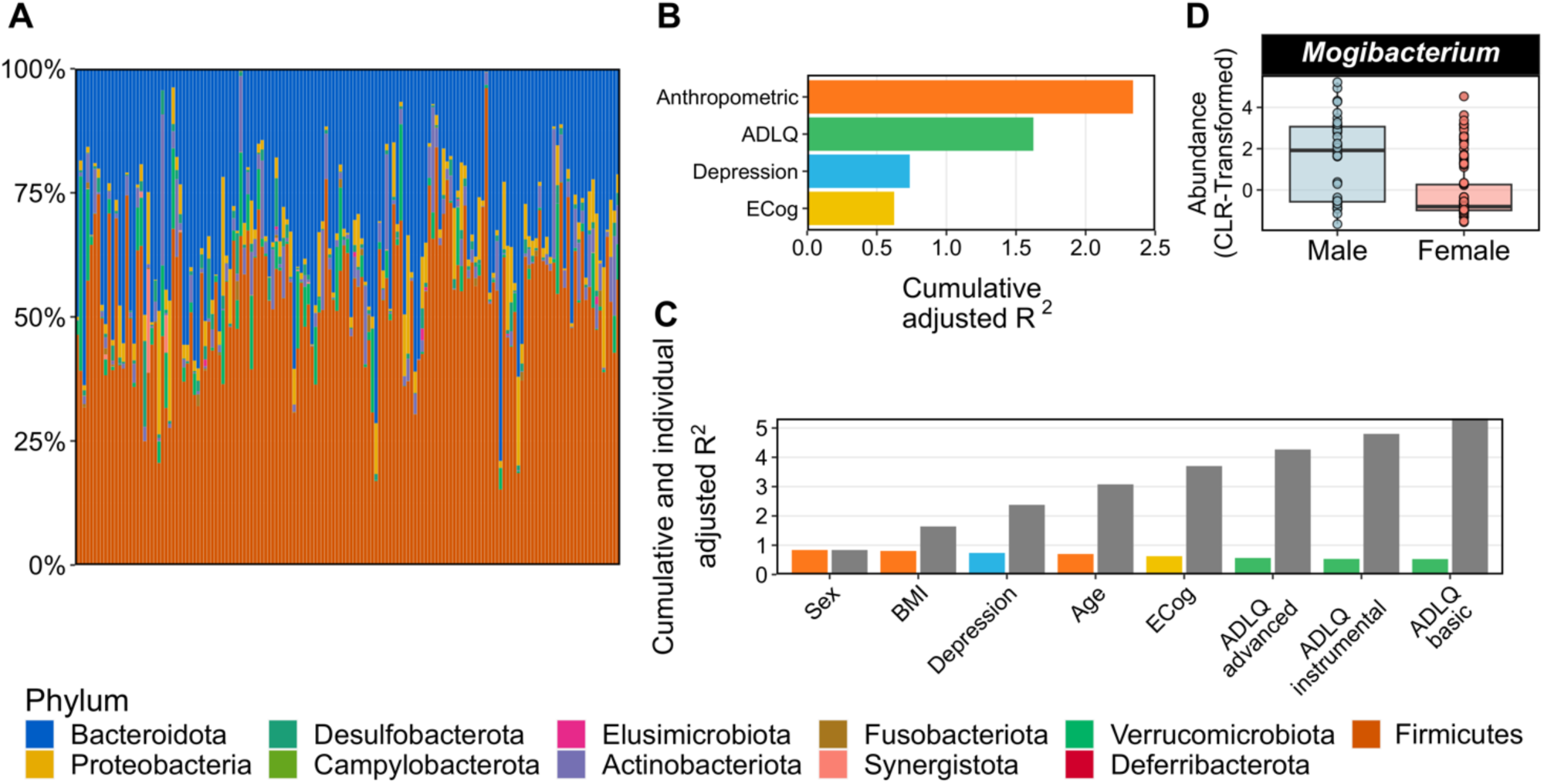
Taxonomic description of the microbiome and explanatory power of cohort covariates on the taxonomic composition of the gut microbiome of the GERO cohort. (A) The relative abundance of the microbiomes for 153 participants of the study. Each colour represents different phyla, as shown in the bottom legend. **(B)** The combined explanatory power of different cohort covariates pooled into 4 pre-defined categories: Anthropometric, ADLQ, Depression and Cognition. Each category is depicted with one colour. **(C)** The individual and cumulative explanatory power of each covariate included in the model used to explain changes in overall microbiome community variation. The colours in the bars correspond to the colours of the 4 categories used in panel B. Gray bars on the right of each coloured bar depict the cumulative effect of the covariates. Coloured bars are sorted in a decreasing order from left to right. **(D)** The genera *Mogibacterium* was found to be more abundant within male participants enrolled in the study. Benjamini-Hochberg method was used to adjust p-values after multiple comparisons. Statistical significance was determined with q < 0.1.

RDA analysis was conducted to disentangle the extent of the overall variation in the bacterial community that could be explained by different variables of the metadata. The anthropometric variables of the participants **(Table 1)** accounted for the greatest amount of variation across the cohort (2.38%), followed by the scores obtained in the ADLQ test (1.56%), their depressive symptoms score according to the GDS (0.74%) and the scores in the ECog test (0.58%) **(Figure 1B)**. We then studied the independent contribution of variables in the metadata by breaking them down. According to the RDA, the three host-derived variables with the most explanatory potential were BMI (0.87%), sex (0.84%) and depressive symptoms score (0.74%), followed by the other anthropometric and cognitive variables, including their ADLQ scores, accounting for a total explained variance of 5.26% **(Figure 1C)**.

**Table 1:** Anthropometric, cognitive and psychological characteristics of the 153 participants aggregated by sex.

| Characteristics aggregated by sex | Female | Male |
| --- | --- | --- |
| Number of participants | 119 (77.77%) | 34 (22.23%) |
| Age | 76.76 ± 5.13 | 76.62±4.69 |
| Height | 152 ± 7 (cm) | 166 ± 7 (cm) |
| Weight | 66.56 ± 12.42 (kg) | 78.31 ± 13.66 (kg) |
| BMI | 28.89 ± 5.28 | 28.16 ± 3.76 |
| Depression Geriatric Scale (GDS) (0-15) | 3.48 ± 3.37 | 2.32 ± 2.41 |
| MOCA/03 | 21.34 ± 4.92 | 21.47 ± 3.16 |
| ACE III (0-100) | 78.55 ± 11.49 | 80.79 ± 9.96 |
| AD8 (0-8) | 2.55 ± 1.78 | 1.91 ± 1.35 |
| ECog | 51.56 ± 14.71 | 50.24 ± 11.42 |
| ADLQ basic | 1.42 ± 4.17 | 0.82 ± 2.29 |
| ADLQ instrumental | 9.17 ± 11.41 | 8.06 ± 7.64 |
| ADLQ advanced | 21.97 ± 18.32 | 27.65 ± 21.35 |
| Dysexecutive questionnaire (DEX) | 10.98 ± 10.63 | 12.19 ± 13 |
| Cognitive Reserve Scale (CRS) | 140.44 ± 26.25 | 132.18 ± 32.88 |

The possibility that the variables in the metadata could explain differences in the abundance of specific bacterial genera was also addressed. The results reveal that the participants’ sex is the only variable that can induce statistically significant changes in the abundance of bacterial genera, with *Mogibacterium* found in higher abundance in males compared to female participants **(Figure 1D)**. Our combined results indicate that although the gut microbiome of the GERO cohort is predominantly composed of the phyla Firmicutes and Bacteroidota, its composition is influenced by anthropometric factors, performance on the ADLQ and ECog tests, as well as depressive symptoms scores.

### 2.2 The gut microbiomes of the GERO cohort show several pathways for the synthesis and degradation of neuroactive molecules

The inferred genomic content from 16S rRNA data was used to infer the abundance of different pathways involved in the synthesis and degradation of neuroactive molecules, the GBMs [29]. Functional analysis revealed the presence of 36 different pathways, representing approximately 64% of the described GBMs [30]. This ratio indicates the existence of different pathways through which the gut microbiomes of the GERO cohort may modulate nervous system function. Additionally, the prevalence of each GBM in the cohort was determined as the percentage of participants in which each step of the GBM was detected as present **(Figure 2A)**. Interestingly, while no GBM was found in less than 5% of the participants, a threshold used to identify rare GBMs, 61% of the GBMs were detected in over 90% of participants, and therefore termed ubiquitous. Particularly, 6 of the 9 GBMs involved in SCFA metabolism (66%) were found in at least 90% of the participants. Additionally, all four GBMs involved in the metabolism of tryptophan and its catabolites were detected in more than 90% of participants. However, pathways involved in the synthesis of important neuroactive molecules such as propionate (a SCFA), and glutamate, the precursor of the neurotransmitter GABA, were present in fewer than 25% of participants. This highlights distinct microbial-neural communication pathways across individuals, potentially providing novel venues for personalised therapeutic and or life-style interventions.

**Figure 2.**
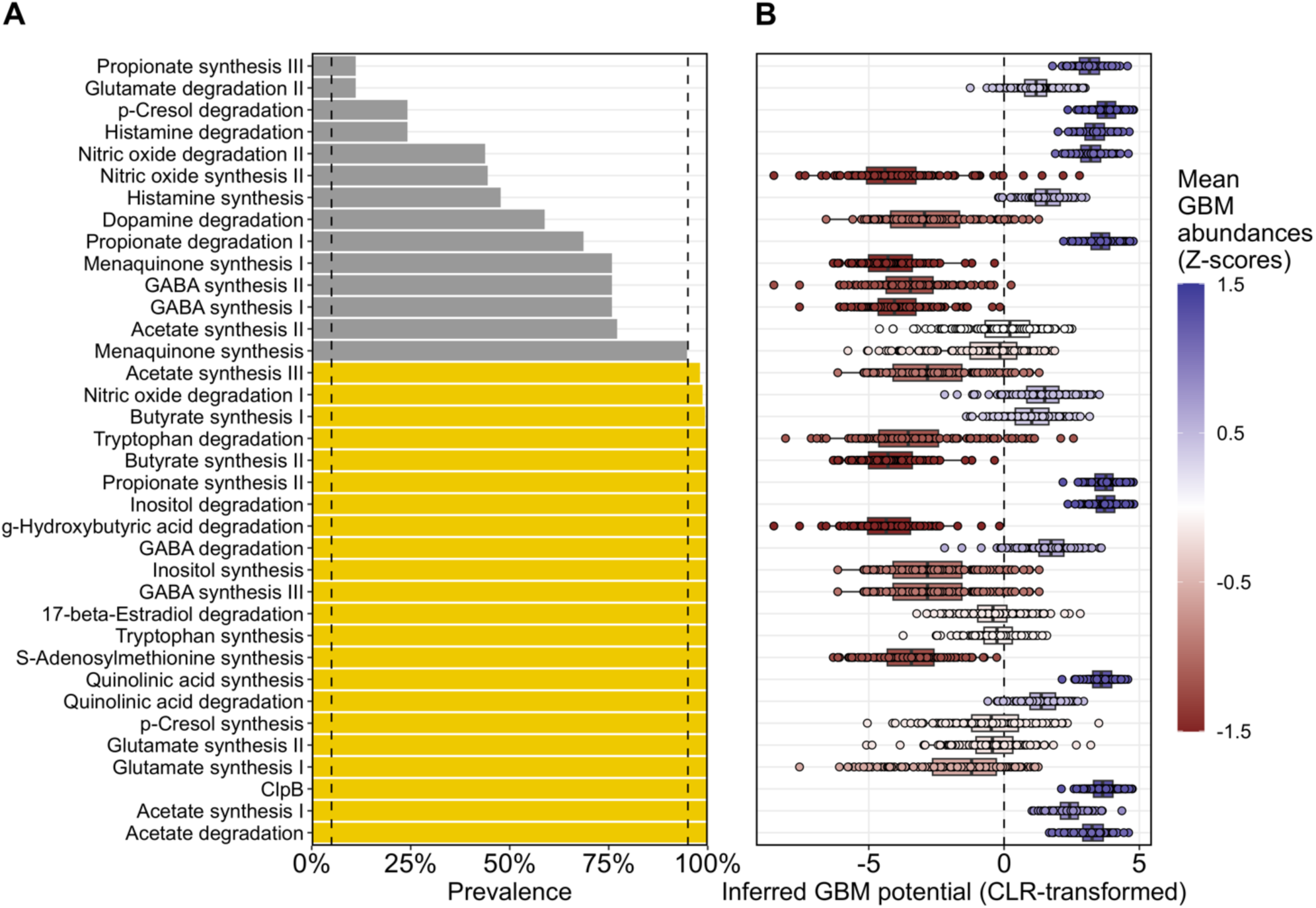
Characterization of the neuroactive potential of the gut microbiome of the GERO cohort. (A) GBMs Prevalence, defined as the proportion of the participants among the total, in which each GBM was fully present. While yellow indicates GBMs with a prevalence above 90%, grey shows GBMs with prevalence ranging from 5% to 90%. **(B)** CLR-transformed abundances of each GBM. Each dot represents one participant. Colours represent Z-scores calculated using the means of the CLR-transformed abundances for all GBMs, with red gradients indicating Z-scores with low abundance (values below 0), and blue gradients indicating Z-scores with high abundance (values above 0).

Additionally, the total abundance of each GBM was calculated for every participant by pooling the metabolic potential of all bacteria in their gut microbiome **(Figure 2B)**. Interestingly, GBMs associated with propionate synthesis, inositol degradation, quinolinic acid synthesis, ClpB, and acetate synthesis and degradation were found in high abundance and prevalence. This may suggest these bacterial pathways play a substantial role in modulating the gut-brain axis in participants of this cohort. On the other hand, GBMs like acetate synthesis, tryptophan degradation, butyrate synthesis, GABA synthesis and degradation, inositol synthesis, and S-adenosylmethionine synthesis were found to be highly prevalent across the cohort but in lower abundances. Conversely, GBMs like propionate synthesis, along with p-cresol, histamine, nitric oxide, and dopamine degradation were found at low prevalence but at high abundances. The high abundance despite their low prevalence may reveal unique neuroactive pathways at the individual level. Our findings demonstrate that the abundance and prevalence of GBMs are not correlated, emphasising the need to consider both aspects when evaluating the neuroactive potential of a microbiome.

### 2.3 Anthropometric variables and ADLQ scores are associated with changes in the neuroactive potential of the gut bacteria

Since we showed that the taxonomic composition of the gut microbiome is influenced by anthropometric and mental variables, we hypothesized that those variables could also modulate the neuroactive potential of the gut microbiome. It was found that the participant scores on the ADLQ basic test, as well as participants’ age and sex have an effect in the abundance of 9 GBMs. Female participants showed a higher potential to synthesise tryptophan, quinolinic acid, p-cresol and glutamate, and also a higher potential to degrade quinolinic acid and acetate. Additionally, our results suggest that as participants’ age increases, their gut microbiome shows a reduced potential to degrade nitric oxide. Our results also showed a decreased potential to synthesise acetate of participants who score higher in the ADLQ basic tests **(Figure 3A)**.

**Figure 3.**
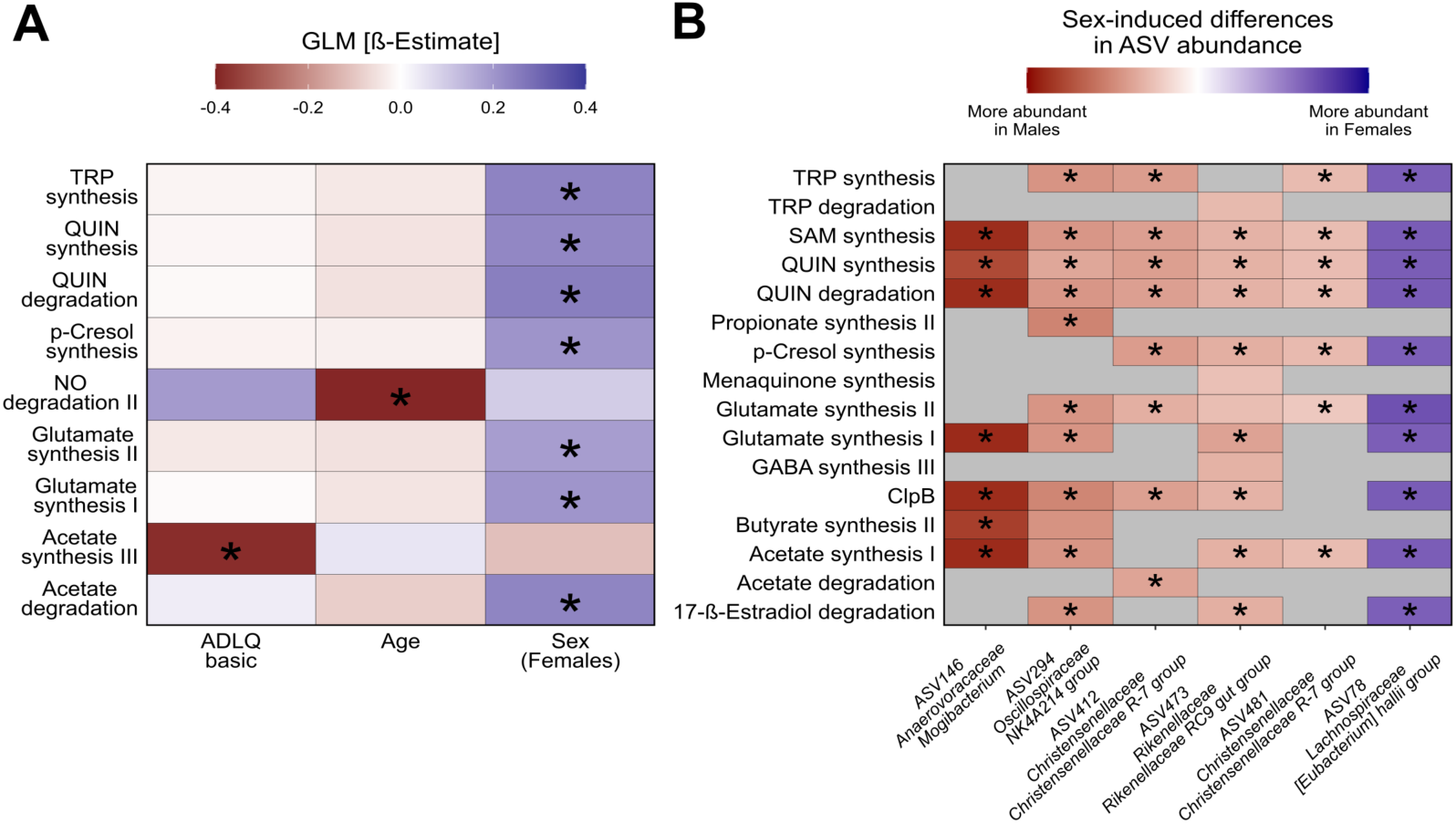
Age, Sex and ADLQ basic test score associated with changes in the neuroactive potential of the gut microbiome. (A) Variations in the overall neuroactive potential of the participant’s gut microbiome associated with changes in their covariates. A higher intensity of blue indicates either a higher abundance of the GBMs as age or ADQL test scores increases, and in female participants. Conversely, a more intense red indicates a lower GMB abundance under the same conditions. **(B)** Sex differences also contributed to statistically significant differences in specific GBMs from six different bacteria. Higher intensities of blue indicate a higher CLR-transformed abundance of the GBM of the specific bacteria in female participants. A more intense red indicates higher CLR-transformed abundances of the GBM in male participants. Grey areas indicate the absence of the GBM in the inferred bacterial genome. Both panels use colour intensity to represent the values of the β estimate in the GLM. Benjamini-Hochberg adjusted p values (q < 0.1) were used for determining significant differences, which are depicted with “*” in the figure panels. Abbreviations: TRP (Tryptophan), QUIN (Quinolinic Acid), NO (Nitric Oxide), SAM (S-Adenosylmethionine).

Additionally, the effects of the same covariates on the neuroactive potential of specific bacteria were tested, but only the sex of the participants was found to induce statistically significant changes. Sex has a significant effect on the neuroactive potential of 6 different ASVs, spanning 5 different genera, altering a total of 16 different GBMs. Results suggest that all the GBMs detected in the genera *Anaerovoracaceae Mogibacterium*, *Christensenellaceae Christensenellaceae R-7 group,* as well as some GBMs detected within the genera *Oscillospiraceae NK4A241* and *Rikenellaceae Rikenellaceae RC9 gut group* are more abundant in male participants. Conversely, all GBMs detected within the *Lachnospiraceae Eubacterium hallii group* were more abundant in female participants **(Figure 3B)**. Together, our results indicate that anthropometric variables such as age and sex, along with the ADLQ basic test scores are associated with changes in the potential to synthesise and degrade neuroactive compounds relevant to the functioning of the CNS.

### 2.4 Depressive symptoms scores and mental test performance are associated with neuroactive potential in the gut microbiome

Several mental tests are used in older adults to evaluate cognitive function and assist in diagnosing neurological and neurodegenerative diseases. It was hypothesised that changes in the gut microbiome composition and its neuroactive potential would be associated with test performance. Our results suggest that higher scores in the AD8 tests are associated with lower gut microbiome potential to degrade dopamine and with higher potential to synthesise butyrate (**Figure 4A)**. On the other hand, higher scores of CRS are associated with a decreased potential of the gut microbiome to synthesise nitric oxide (**Figure 4A)**. Although functional changes in the gut microbiome were associated with participant score in two mental tests, no statistically significant associations were found between the abundance of any gut-bacterial taxa and test scores.

**Figure 4.**
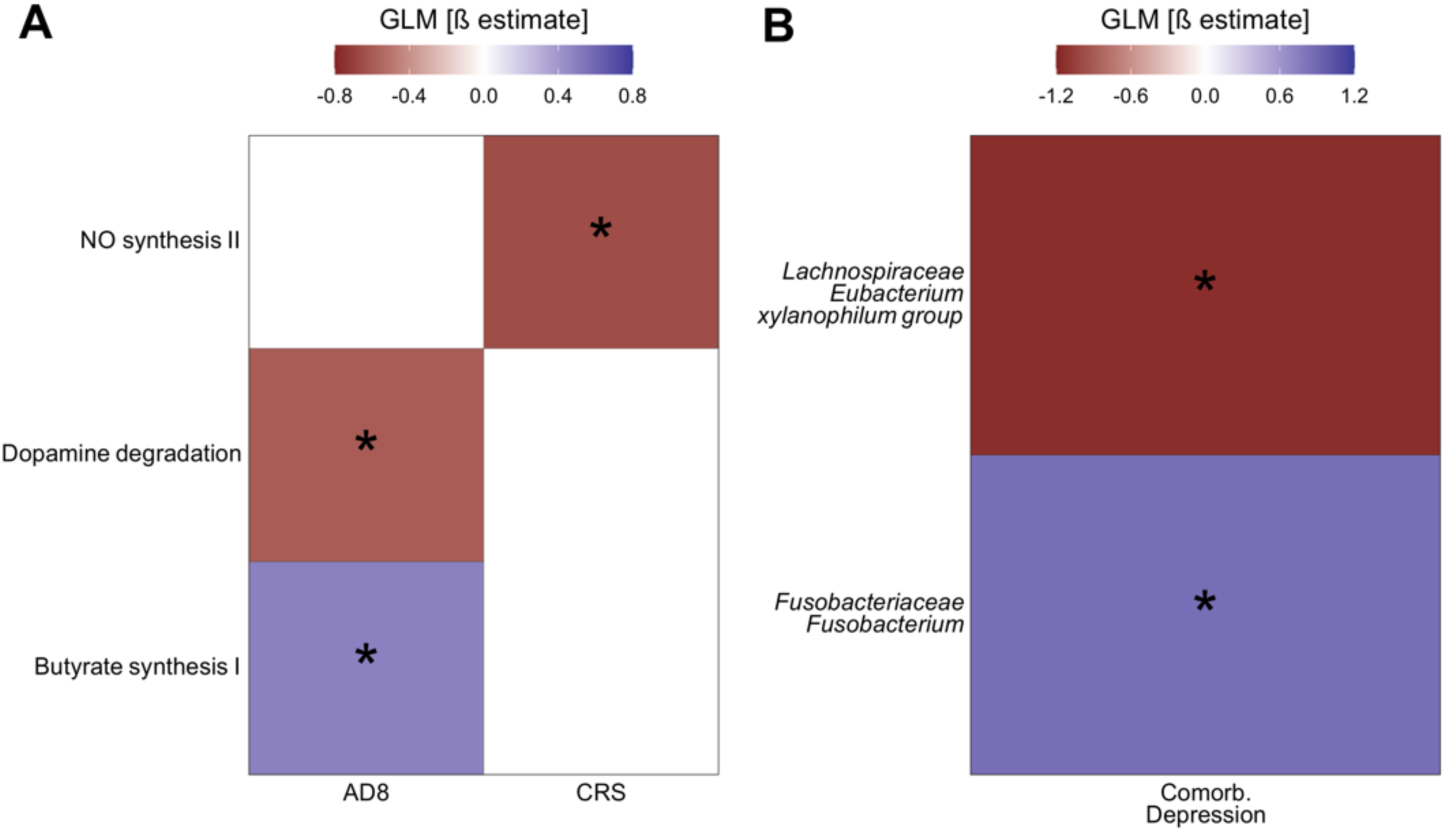
Mental test scores are associated with changes in the neuroactive potential of the gut microbiome and depressive symptoms are linked to specific gut bacterial taxa. (A) Associations between scores in the mental tests AD8 and CRS and the neuroactive potential of the gut microbiome. While blue represents a higher GBM abundance in participants with higher scores in the tests, red represents the opposite. **(B)** Associations between the abundance of genera and the depressive symptoms of the participant. “Comorb. Depression” denotes patients with depressive symptoms as a comorbidity. Blue represents a higher abundance of the bacteria in depressive participants, and red represents the opposite. In both panels, colours represent the β estimate of the GLM. Benjamini-Hochberg adjusted p values (q < 0.1) were used for determining significant differences and are depicted with “*” in the figure panels.

Additionally, it was evaluated whether participants self-reporting depression showed distinct gut microbiome compositions or different neuroactive potentials when compared to the other participants. Results indicate that participants with self-reported depression harbour a decreased abundance of the genera *Lachnospiraceae Eubacterium xylanophilum group*, and a higher abundance of the genera *Fusobacteriaceae Fusobacterium* (**Figure 4B**). However, no statistically significant changes were found in the neuroactive potential of their gut microbiomes. Together, these results link AD8 and CRS test scores to specific changes in the metabolism of nitric oxide, dopamine, and butyrate, which are not associated with the abundance of specific bacteria. On the other hand, in participants with depressive symptoms, two bacteria from the gut microbiota were found to be associated with this comorbidity.

## 3. Discussion

There is a growing emphasis on the potential of the microbiome to be a hallmark of aging in health and disease [26]. To the best of our knowledge, this is the first time the composition and neuroactive potential of the gut microbiomes of a population of elderly South Americans with cognitive complaints is associated with their performance in tests evaluating participants’ mental health and functional decline. Specifically, we found changes in the abundance of two bacterial genera in participants with self-reported depression. Additionally, changes in the abundance of functional pathways involved in the metabolism of neuroactive compounds were associated with changes in participants’ score in the AD8 and CRS tests. Therefore, our findings provide new insights into the role of intestinal microbiota in the cognitive decline and mental health of an ageing population with cognitive complaints from the global south.

The gut microbiome of the cohort shows a taxonomic composition **(Figure 1A)** similar to other human populations, dominated by Firmicutes and Bacteroidetes, followed by less predominant phyla such as Actinobacteria, Proteobacteria and Verrucomicrobiota [42,43]. Previous research on a younger and smaller group of Chilean participants showed similar findings [44]. One noticeable difference however pertains to the genera *Akkermansia*. Here, as well as in the younger cohort of Chileans, *Akkermansia* was the only genera within the phylum Verrucomicrobiota (previously known as Verrucomicrobia) detected. However, while it was previously identified as part of the core Chilean microbiota (defined in that article as genera that were present in every individual of the cohort), in the GERO cohort, sequences assigned to *Akkermansia* were found only in 66.0% of the participants. Although evidence has shown that *Akkermansia* is more abundant among the healthy aged populations [26,28], the evidence regarding changes in its prevalence is less convincing.

Our analysis confirms that sex, age, and BMI significantly, albeit moderately, influence the taxonomic composition of the gut microbiome **(Figure 1B,C)**, similar to other European cohorts [30]. Additionally, it was previously shown that non-anthropometric variables like the quality of life of a participant, assessed by using the RAND 36-item survey, could explain up to 1.48% of the overall taxonomic changes in the gut microbiome [30]. Interestingly, we report a close percentage (1.56%) of explained taxonomic variation using the three ADLQ questionnaires **(Figure 1B)**, which evaluate participants’ functional capacity. Although developed with a different scope, ADLQ assesses participants’ quality of life, physical functioning and mental health. Therefore, our results using ADLQ align with previous observations of the effects of the participants’ quality of life on shaping the gut microbiome composition [30].

Interestingly, it has been reported that men perform worse than women in memory, mental speed and global cognitive function in an elderly cohort [45], and that patients with cognitive complaints showed an increased abundance of *Mogibacterium* [46,47]. Our results showed a higher abundance of the genus *Mogibacterium* among male participants within a cohort of participants reporting cognitive complaints **(Figure 1D)**, supporting the observations of the aforementioned works. This may implicate a role of the gut microbiome regarding sex differences in cognitive function and healthy ageing in humans.

The prevalence of GBMs within the gut microbiome of the participants was assessed. Our results align with a previous observation [30], as we found that all the GBMs previously defined as ubiquitous within bacterial genomes were present in more than 90% of the cohort participants, and GBMs previously defined as rare were also identified as less prevalent in the gut microbiome of the participants **(Figure 2A)**. Interestingly, our results suggest that the extent of the prevalence of a GBM is not indicative of how abundant that GBM is within the bacterial community **(Figure 2B)**. However, the biological interpretation of these observations remains elusive, and more research is warranted.

Our results suggest a consistent effect of participants’ sex on the potential of their gut microbiome to metabolise tryptophan (TRP) and its derived metabolite quinolinic acid (QUIN) **(Figure 3A)**. On the one hand, TRP has been implicated in the modulation of the gut barrier permeability, either directly [48], or through catabolites generated by the metabolic activity of the gut microbiome [49]. Moreover, subsequent changes in permeability could in turn alter the exchange of TRP, QUIN and other metabolites between the gut lumen and the bloodstream, from where they can reach the brain. For instance, TRP can reach the brain after its transportation through carriers [50]. Conversely, QUIN, a molecule with neurotoxic properties, has been associated with neurological disturbances that can become more prevalent as individuals grow older [51], has been shown to accumulate in animal models as they age, particularly in the context of inflammatory neurological diseases [52,53]. Additionally, an increase in QUIN level in blood has been linked to increased prevalence of Alzheimer’s disease [54,55], which is more prevalent in women than men [56,57].

Our results also suggested that female participants showed an increased potential to synthesise p-cresol **(Figure 3A)**, a molecule whose abundance in the stool of aged humans has been linked with increased frailty in the ELDERMET cohort [58]. Additionally, another study integrating three independent cohorts showed that p-cresol and its associated metabolites were found in higher abundances in aged participants, where an effect of the sex of the participant on the abundance of the molecule was also described [17]. This further highlights the relevance of our findings showing sex-induced differences in the gut microbiome potential to synthesise p-cresol and possible implications in different ageing trajectories between the two biological sexes.

Our studies reveal that the gut microbiomes of female participants exhibited an increased potential to degrade acetate and that a reduced potential of the gut microbiome to synthesise acetate is associated with higher scores in the ADLQ basic test, an indicator of a higher level of dependence **(Figure 3A)**. Acetate levels decrease in aged mice, and its supplementation improved their health span along ageing [59]. Moreover, other work shows how the removal of acetate-producing bacteria from the gut of aged mice is associated with learning and memory complaints, along with synaptophysin reduction in the hippocampus [60], a protein important for synapse formation within the hippocampus [61]. Interestingly, mice synaptophysin levels could be restored after acetate supplementation [60]. While additional work is necessary to understand the translational implications of these findings, the evidence highlights the relevance of our results as it emphasises sex-driven differences in the metabolism of a molecule crucial to healthy ageing in animal models.

Additionally, we found that as the participants’ age increases, the gut microbiome potential to degrade NO decreases **(Figure 3A)**. This may suggest an accumulation of NO in the gut environment, which has been linked to increased gut permeability [62], a condition usually associated with health challenges in older individuals [25,63]. Nevertheless, more studies are needed to further investigate the role of the gut microbiome in aged cohorts from the global south. In that respect, other experimental designs such as cross-sectional or longitudinal studies may unveil further insights into the role of the gut microbiome in healthy ageing in South American subjects.

We also reported associations between the participants’ performance on the AD8 and CRS tests, and the neuroactive potential of their gut microbiome, which are the first of their kind described in populations outside the global north. As mentioned before, the AD8 test is an informant-based questionnaire aimed at early detection of cognitive changes indicative of dementia or cognitive decline associated with Alzheimer’s disease [12]. Interestingly, a meta-analysis of cross-sectional studies suggests that increased inflammation is associated with Alzheimer’s disease and mild cognitive impairment among the elderly [64]. Noteworthily, butyrate [65] and dopamine [66] in the gut have been implicated in the modulation of systemic inflammation, which aligns with our results showing associations between the scores in AD8 and both metabolites **(Figure 4A)**. This suggests a potential indirect mechanism by which the gut microbiome may influence the cognitive performance of the elderly through the modulation of host systemic inflammation, a concept supported by existing research linking the gut microbiome to the modulation of this process [26].

Additionally, we observed that higher scores in the CRS test were associated with a reduced potential of the gut microbiome to degrade NO **(Figure 4A)**. Importantly, NO has been implicated in regulating gut permeability [62], and modulating the synthesis of brain-derived neurotrophic factor (BDNF) in the brain [67]. BDNF plays an important role in neuronal survival, growth, and neural plasticity [68]. Interestingly, neural plasticity is a key mechanism in cognitive reserve and a pivotal characteristic that accounts for individual differences in susceptibility to age-related brain changes or Alzheimer’s disease-related pathology [69]. However, further research should explore how alterations in intestinal NO levels might affect the production of BDNF in the brain, and its potential to cross the brain barrier. Additionally, we found that participants with self-reported depressive symptoms exhibited a lower abundance of the genera *Lachnospiraceae Eubacterium xylanophilum group*, and a higher abundance of the genera *Fusobacteriaceae fusobacterium* **(Figure 4B)**. Interestingly, there is still a lack of consensus regarding the association of specific bacteria and depression symptoms. While higher abundance of *fusobacterium* in patients with depressive symptoms has been shown [70], the opposite association has also been reported [71]. To our knowledge, there have been no previous reports of an association between the *Lachnospiraceae Eubacterium xylanophilum group* and depression, although a member of the *Lachnospiraceae* family was previously associated with depressive symptoms [72]. Noteworthy, associations between abundances of bacteria and symptoms or self-reported illnesses must be taken with care, especially when the gut microbiome is profiled with marker genes [73].

It is worth noting that this study is not without its limitations. Firstly, the absence of a group of elderly individuals without cognitive dysfunction used as a control makes this an observational study. Additionally, it remains to be understood to what degree the observed mood effects are specific to cognitive dysfunction and whether they can be generalized to a cognitive decline-free population. However, a closer inspection of experimental designs used in clinical studies have shown that the magnitude of the effects of treatment in well-designed observational studies don’t systematically differ with those reported in experimental designs of higher levels of evidence [74]. Additionally, the plethora of lifestyle patterns known to modulate the composition of the gut microbiome, such as diet, physical activity, accessibility to green areas and exposure to xenobiotics and environmental pollutants couldn’t be totally accounted for in this study. Future investigations, especially emerging from the Global South, should address these factors [75]. Finally, besides the aspects already discussed, future studies conducted in similar populations should also consider the temporal dynamics of the human gut microbiome for a personalised understanding of health- and disease-associated microbiome patterns [76].

In conclusion, this study has characterised the gut microbiome composition and neuroactive potential of a population of elderly Chileans, as well as described its significant association with participant’s anthropometric variables, scores in mental tests, and depression. Particularly, participant’s gut microbiome potential to metabolise TRP, SCFAs, p-cresol, glutamate, and NO was found to vary with either age and sex, and performance on mental tests such as the basic ADLQ, AD8, and CRS. Moreover, two bacterial genera -*Lachnospiraceae Eubacterium xylanophilum group* and *Fusobacteriaceae Fusobacterium*-showed altered abundances in patients who self-reported depression. Interestingly, this work consistently highlights that the neuroactive potential of gut microbiota in the elderly with cognitive complaints varies between men and women, which could be relevant to deepen our knowledge about other sex-specific differences produced during ageing. However, further studies are required to fully understand the role of the gut microbiome in the ageing brain.

## 4. Materials and methods

### 4.1 Cohort

The population of 153 individuals described in this study corresponds to the baseline assessment of the Chilean GERO cohort. A community-based cohort aimed at analysing predictors of functional decline and dementia progression in elderly individuals with cognitive complaints residing in the community [9]. This is a clinical project of the Geroscience Center for Brain Health and Metabolism. Cohort participants were recruited from the general population using a door-to-door strategy between November 2017 and December 2021 in Santiago, Chile. Participants were required to be 70 years or older with cognitive complaints reported by themselves or a reliable informant. Additionally, participants needed to have a Clinical Dementia Rating Scale (CDR-FTLD) score equal to or less than 0.5, live in the community, and have a knowledgeable informant. Exclusion criteria included diagnosed dementia or screening tests indicating dementia, cognitive impairment (Mini-Mental State Examination (MMSE) scores < 21) [77], and functional impairment (Pfeffer questionnaire score > 2) [78]. Additionally, participants were excluded if they had a history of institutionalisation, illiteracy, severe sensory or mobility impairments, major psychiatric or neurological disorder (such as schizophrenia, brain tumour, subdural hematoma, progressive supranuclear palsy, head trauma, or recent stroke), or a fatal with a prognosis of less than one year. All study participants had a reliable informant who provided information about their functional abilities. After inclusion, all subjects completed an extensive neurological, neuropsychological, and neuroimaging evaluation (see Slachevsky, et al., 2020 [9] for a detailed description of the protocol). Each participant provided signed informed consent, and the study was approved by the Ethic Committee of the Servicio de Salud Metropolitano Oriente, Santiago (Chile). The project is funded by the Chilean National Agency of Research (ANID, FONDAP 15150012).

### 4.2 Data collection

Anthropometric measurements, blood tests, and stool collection, as well as neuropsychological assessments involving cognitive, functional, and neuropsychiatric tests, were conducted [9]. The results of these questionnaires were collected concurrently with the biological samples, ensuring that the data from both sources were gathered during the same period. Anthropometric data included age, sex and BMI. Cognitive status was assessed using the Montreal Cognitive Assessment (MoCA) and the Addenbrooke’s Cognitive Examination III (ACE-III), covering five cognitive domains [79]. The MoCA score ranges from 0 to 30, with higher scores indicating better cognitive performance, while the ACE-III scores range from 0 to 100. Exclusion criteria include psychiatric or neurological disorders, with severe depression participants not being part of the cohort. However, depressive symptoms were assessed using the Geriatric Depression Scale (GDS), with scores ranging from 0 to 15 [77,78,80–82]. Informant-rated questionnaires, such as the Technology-Activities of Daily Living Questionnaire (T-ADLQ) were used to assess everyday functioning in three domains: Basic, Instrumental, and Advanced. The measurement range for each domain is from 0 to 100%, with higher percentages indicating greater functional impairment [10,83,84]. Additionally, the Everyday Cognition (ECog) questionnaire was used to measure cognitive abilities across six domains with a total of 39 questions, each scored from 1 to 5, with a higher total scores indicating more impairment [85]. Executive dysfunction was assessed using the Dysexecutive Questionnaire (DEX) test (range from 0 to 80, where the number indicates the degree of executive dysfunction), and the AD-8 questionnaire, consisting of 8 questions with a total score of 8, was used to evaluate functional changes typically associated with dementia [12,13]. Finally, the participation in cognitively stimulating activities across different life stages was assessed using the Cognitive Reserve Scale (CRS) test, with a scale from 0 to 15 [14,86]. It is important to mention that the depressive status of participants was determined through self-reporting according to the guidelines in the National Health Survey [87].

### 4.3 Sample collection

Samples were obtained at the Memory and Neuropsychiatric Clinic (CMYN) Neurology Department, Hospital del Salvador and Faculty of Medicine, University of Chile, Santiago, Chile [9]. The stool samples were collected in OMNIgene gut tubes (DNAgenotek, OMR-200), and the participants performed the sample collection at home following the instructions provided by a nurse. The tubes containing the samples were then retrieved and stored for no more than one month at room temperature until their arrival at the laboratory, where they were kept at -80 °C until processing.

### 4.4 Sample processing

DNA extraction from the samples was carried out utilising the IHMS SOP 06 V1: STANDARD OPERATING PROTOCOL FOR FECAL SAMPLES (human-

microbiome.org/index.php?id=Sop&num=006) and QIAamp Fast DNA Stool Mini Kit (Quiagen, 51604). The extraction process, as well as quality control, were conducted at the Center for Integrative Biology, Mayor University. A gut microbiome biobank was established and preserved at -80 °C. The DNA samples were transported on dry ice for sequencing.

### 4.5 Sequencing

DNA sequencing was performed at the Alkek Center for Metagenomics and Microbiome Research (Department of Molecular Virology and Microbiology at Baylor College of Medicine, Houston, Texas, USA). The V4 region of the 16S rRNA gene was targeted using the primers 515F: GTGCCAGCMGCCGCGGTAA and 806R: GGACTACHVGGGTWTCTAAT [87]. The sequencing procedure involved performing 2x250 bp paired-end sequencing using an Illumina Miseq system in rapid run mode.

### 4.6 Bioinformatics and statistical analysis

#### 4.6.1 16S data analysis, taxonomic, functional annotation, and diversity within samples

FastQ files with raw sequences were loaded into R (version 4.1.1), where the Divisive Amplicon Denoising Algorithm (DADA2) v. 1.22.0 [88] was used to construct a count table of amplicon sequence variants (ASV). The DADA2 workflow consisted of filtering and trimming low-quality reads, learning error rates, sample inference, merging paired reads, constructing an ASV table, removing chimeras and assigning taxonomy up to the genus level. For each step of the workflow default parameters were used with the exception of the function filterAndTrim from the dada2 R package. The following non-default values were used as arguments of the function: maxEE was set as c(2,5), truncQ as 2, trimLeft as 20 and truncLen as 200 to fix the length of the sequences used later in the analysis. Then, for taxonomy assignment, SILVA database (v 138) was used as a reference [89]. PICRUSt2 (version 2.4.1) [90] was used with its default parameters to infer the genomic content from 16S data, using the KEGG database [91] as a reference. The inferred genomic content was used to compute the abundance of Gut Brain Modules (GBMs), defined as a collection of 56 bacterial biochemical pathways involved in the synthesis and degradation of neuroactive molecules [30]. To determine the abundance of the GBMs, the OmixerRpm software [92] was used in stratified mode (i.e., determine GBMs for each independent bacterial genera), and in the unstratified mode (i.e., the combined profiles of every bacterium from a participant’s microbiome). In both analyses, default software parameters were used. For following analyses, the ASV table and the GBMs table were transformed using the Centered Log Ratios (CLR) transformation implemented in the vegan package with a pseudocount of 2/3. Downstream statistical analysis was performed with R (version 4.2.0) and RStudio (version 2022.7.1.554).

#### 4.6.2 Microbiota community variation explained by metadata variables

Variables included in our analyses were first checked to avoid high collinearity before statistical models were produced, for which a Pearson’s rho value higher than 0.7 was used as an exclusion cutoff. Variables in the metadata were scaled before being included in the models. First, redundancy analysis (RDA) was used to determine the contribution of variables in the metadata to the variations observed in the taxonomic composition of the gut bacteria among participants. RDA was performed using the vegdist function from the vegan package (version 2.6.4). Dissimilarity was calculated using Aitchison’s distance (Euclidean distance on CLR-transformed data) as recommended for the analysis of compositional data [93]. For the analysis, variables previously found to influence the taxonomic composition of the gut microbiome were included in our statistical models. Therefore, we aimed to explain differences in the overall composition of the gut microbiome (i.e., the complete CLR-transformed bacterial counts matrix) using the following formula: “matrix ∼ sex + age + bmi + self-reported depression + cognition + ADLQbasic + ADLQinstrumental + ADLQadvanced + batch”. The last variable, batch, was included to account for possible batch effects due to different sequencing runs. The same covariates were used to identify possible differences in the abundances of specific bacterial genera. To do so, Generalized Linear Models (GLMs) were fitted using the same formula used for the RDA analysis with the function fw_glm from the Tjazi R package, in accordance recent recommendations [94,95]. Calculated p-values were adjusted using the FDR Benjamini-Hochberg method. Adjusted p-values < 0.1 were deemed as statistically significant.

#### 4.6.3 Description of the neuroactive potential of the GERO cohort

With the count tables of GBMs abundances and coverages, the prevalence of each GBM in the cohort was calculated as the percentage of the total of participants in which the GBM was detected. Importantly, GBMs were deemed present in the gut microbiome of a patient only if all the genes that were originally described as part of each GBM [30] were inferred to be present, in order to ensure a stringent detection criterion. Additionally, to assess if variables previously found to influence the taxonomic composition of the gut microbiome could also influence its neuroactive potential, a GLM for each GBM was fitted using the following formula “GBM_abundance ∼ sex + age + bmi + GDS total score (ts) + ECog ts + ADLQbasic ts + ADLQinstrumental ts + ADLQadvanced ts + batch” as required by the function fw_glm from the Tjazi R package [94,95]. The same formula was used then to assess if those variables could also induce changes in the abundance of GBMs encoded in specific bacterial genera (i.e., using the stratified GBMs count table produced with omixerRpm software as explained previously). In both cases, the variable batch was included to account for possible batch effects due to different sequencing runs, and the calculated p-values were adjusted using the FDR Benjamini-Hochberg method. The adjusted p-values < 0.1 were deemed as statistically significant.

#### 4.6.4 Associations between participants’ scores in mental tests and their gut microbiome

Scores achieved by participants in a battery of mental tests were used to test three hypotheses, namely: that (1) the taxonomic composition of the gut microbiome, or (2) the neuroactive potential of certain bacterial genera, or (3) the neuroactive potential of the entire gut microbiome are associated with the participant’s score in the mental questionnaires. To test these hypotheses, linear models were fitted using the scores of the participants in different mental tests as covariates explaining the abundance of either (1) a bacterial genus, (2) each GBMs found within each bacterial genus or (3) the abundance of each GBM in the entire gut microbiome. For the three hypotheses, the fw_glm function from the Tjazi package was used to fit GLMs, and the formula included the total scores (ts) of all the mental tests each participant answered as follows: “response ∼ AD8 ts + MoCA ts + DEX ts + ECog ts + ACEIII ts + CRS ts + self-reported depression + batch”. The variable batch was included to account for possible batch effects due to different sequencing runs. Calculated p-values were adjusted using the FDR Benjamini-Hochberg method, and adjusted p-values < 0.1 were deemed as statistically significant.

## 4.7 Data availability

The raw sequences generated for this project were deposited in the ENA and are publicly available under the accession number PRJEB77146.

## 5. Abbreviations

ACE-III: Addenbrooke’s Cognitive Examination III
AD8: Alzheimer Disease 8
ADLQ: Activities of Daily Living Questionnaire
ASV: Amplicon Sequence Variants
BDNF: Brain-Derived Neurotrophic Factor
BMI: Body Mass Index
CDR: Clinical Dementia Rating Scale
CLR: Cantered Log Ratios
CRS: Cognitive Reserve Scale
DEX: Dysexecutive Questionnaire
ECog: Everyday Cognition
ENA: European Nucleotide Archive
FDR: False Discovery Rate
GDS: Geriatric Depression Scale
GBM: Gut Brain Modules
GLM: Generalized Linear Model
MMSE: Mini-Mental State Examination
MoCA: Montreal Cognitive Assessment
NO: Nitric Oxide
QUIN: Quinolinic Acid
RDA: Redundancy Analysis (RDA)
SAM: S-adenosylmethionine
SCFA: Short Chain Fatty Acids (SCFA)
T-ADLQ: Technology-Activities of Daily Living Questionnaire
TRP: Tryptophan

## 6. Author Contributions

PCR and BV drafted the manuscript. AS, CGB, and FC designed the study. PCR conducted the laboratory processing of the samples. BV performed the bioinformatic and statistical analyses and prepared the figures. All authors revised and edited the manuscript. The final version was approved by all authors.

## 8. Ethical Statement

Each participant provided signed informed consent, and the study was approved by the Ethic Committee of the Servicio de Salud Metropolitano Oriente, Santiago (Chile).

## 9. Founding

This study was supported by the “*Fondo de Financiamiento de Centros de Investigación en Áreas Prioritarias*” ANID/FONDAP/15150012 (Chile).

